# Assemblies of long-read metagenomes suffer from diverse errors

**DOI:** 10.1101/2025.04.22.649783

**Authors:** Florian Trigodet, Rohan Sachdeva, Jillian F. Banfield, A. Murat Eren

## Abstract

Genomes from metagenomes have revolutionised our understanding of microbial diversity, ecology, and evolution, propelling advances in basic science, biomedicine, and biotechnology. Assembly algorithms that take advantage of increasingly available long-read sequencing technologies bring the recovery of complete genomes directly from metagenomes within reach. However, assessing the accuracy of the assembled long reads, especially from complex environments that often include poorly studied organisms, poses remarkable challenges. Here we show that erroneous reporting is pervasive among long-read assemblers and can take many forms, including multi-domain chimeras, prematurely circularized sequences, haplotyping errors, excessive repeats, and phantom sequences. Our study highlights the need for rigorous evaluation of the algorithms while they are in development, and options for users who may opt for more accurate reads than shorter runtimes.

## Introduction

By enabling massively parallel sequencing of DNA molecules, the second generation sequencing technologies that emerged in the early 2000s had a remarkable impact on all disciplines of life sciences (Shendure and Ji 2008; van Dijk et al. 2014). A crucial advancement brought by this technology in microbiology was the ability to reconstruct microbial genomes directly from environmental ‘metagenomes’ without cultivation (Tyson et al. 2004), which substantially enhanced our understanding of microbial diversity and function and continues to be an indispensable tool for biotechnology and biomedicine (Eren and Banfield 2024). Yet, the relatively short sequencing reads produced by the second generation sequencing posed significant limitations on assembly (Olson et al. 2019; Tørresen et al. 2019), a critical computational step in genome recovery workflows where reads are stitched together to rebuild contiguous segments of DNA (contigs) prior to binning, and often led to highly fragmented and sometimes highly contaminated genomes from metagenomes (Shaiber and Eren 2019; Chang et al. 2024; Chen et al. 2020).

The emergence of the third generation sequencing technologies, such as those implemented by Pacific Biosciences (PacBio) and Oxford Nanopore Technologies (ONT), marks another breakthrough in genome-resolved metagenomics through ultra-long (Jain et al. 2018) and increasingly accurate reads that can help solve complex genomic puzzles with unprecedented precision (Wenger et al. 2019). These new opportunities rely heavily on the advancement of long-read assembly algorithms, an area of active research with multiple successful software products that can assemble long-reads into complete chromosomes (Feng et al. 2022; Benoit et al. 2024; Kolmogorov et al. 2020; Nurk et al. 2020). However, assessing the accuracy of the assembled long reads poses remarkable challenges, especially from complex environments that often include poorly studied organisms but host substantial novel diversity.

The benchmarking of assembly algorithms typically relies on a few key principles, including the use of mock or simulated datasets, evaluation of contig length distributions, assessment of unassembled read fractions, genes, functions, and comparisons with other assemblers (Mikheenko, Saveliev, and Gurevich 2016; Meyer et al. 2022; Delgado and Andersson 2022). While these strategies are critical for algorithm development, benchmarking efforts rarely take into account how individual reads align to assemblies and quantify the rate, origins, and impact of mismatches between original reads and final contigs. Here we assessed the extent of agreement between individual high-fidelity PacBio reads and assembly results reported by four state-of-the-art long-read assemblers, and observed a wide range of issues, including haplotyping errors, chimerism, premature circularization, and regions of contigs that are not supported by an of the input sequences. Based on our prior experience (e.g., (Schoelmerich et al. 2024)), assemblies of ONT long reads exhibit similar errors, yet we limit our benchmarks to PacBio given the relatively higher base accuracy of PacBio reads (∼99.95%, or 5 errors per 10 kbp) compared to the ONT reads (∼99%, or 100 errors per 10 kbp). Overall, our survey offers reproducible means to identify long-read assembly errors and insights into their downstream implications for researchers who develop or use long-read assembly algorithms to consider.

## Results

Our study benchmarks four state-of-the-art long-read assembly software, (1) HiCanu v2.2 (Nurk et al. 2020), (2) hifiasm-meta v0.3 (Feng et al. 2022), (3) metaFlye v2.9.5 (Kolmogorov et al. 2020) and (4) metaMDBG v1 (Benoit et al. 2024), based on their assembly performance of 21 PacBio HiFi metagenomes (Supplementary Table 1). Thirteen of these metagenomes were used by the authors of at least one assembly algorithm to evaluate performance. They include mock communities and anaerobic digesters, as well as gut samples from humans, chickens, and sheep.

The remaining eight are novel HiFi metagenomes from the surface ocean, a key biome that was not included in previous benchmarks and represents a complex ecosystem with little genomic representation (Schechter et al. 2025) (Supplementary Table 1).

Investigations of the agreement between individual long reads and assemblies they support require the alignment of individual reads to resulting contigs. For this task our study employs minmap2, a popular and high-performance mapping software designed to align long-reads to reference sequences (Li 2018). Unlike short-read alignment software such as Bowtie2 (Langmead and Salzberg 2012) or BWA (Li and Durbin 2009), minimap2 allows premature ends in read alignments and can map the remainder of the read to another reference location. This so-called ‘read clipping’ procedure is a critical tool to identify locations in the final assembly that is poorly supported by individual long-reads. To gain quantitative insights into assembly artifacts we developed ‘<u>anvi-script-find-misassemblies</u>’, a script that comprehensively summarizes locations and frequencies of read clipping events, where long reads are split systematically by the mapping software to maximize agreement between reads and assemblies, in addition to regions in contigs that are not supported by individual long reads.

### Assembly errors are common to all long-read assemblers

Instances where the vast majority of long-reads are clipped at the same nucleotide position is strong evidence that the final assembled sequence contains an error. However, quantifying the number of errors in a given assembly is not as straightforward as counting the clipping events since a single assembly error may result in multiple clipping events associated with neighboring nucleotide positions in the reference sequence (Figure 1A). To minimize false positives in our results, and exclude clipping events due to within-population biological variation, our analyses below primarily focus on locations in long-read assemblies with 100% clipping rate.

**Figure 1.**
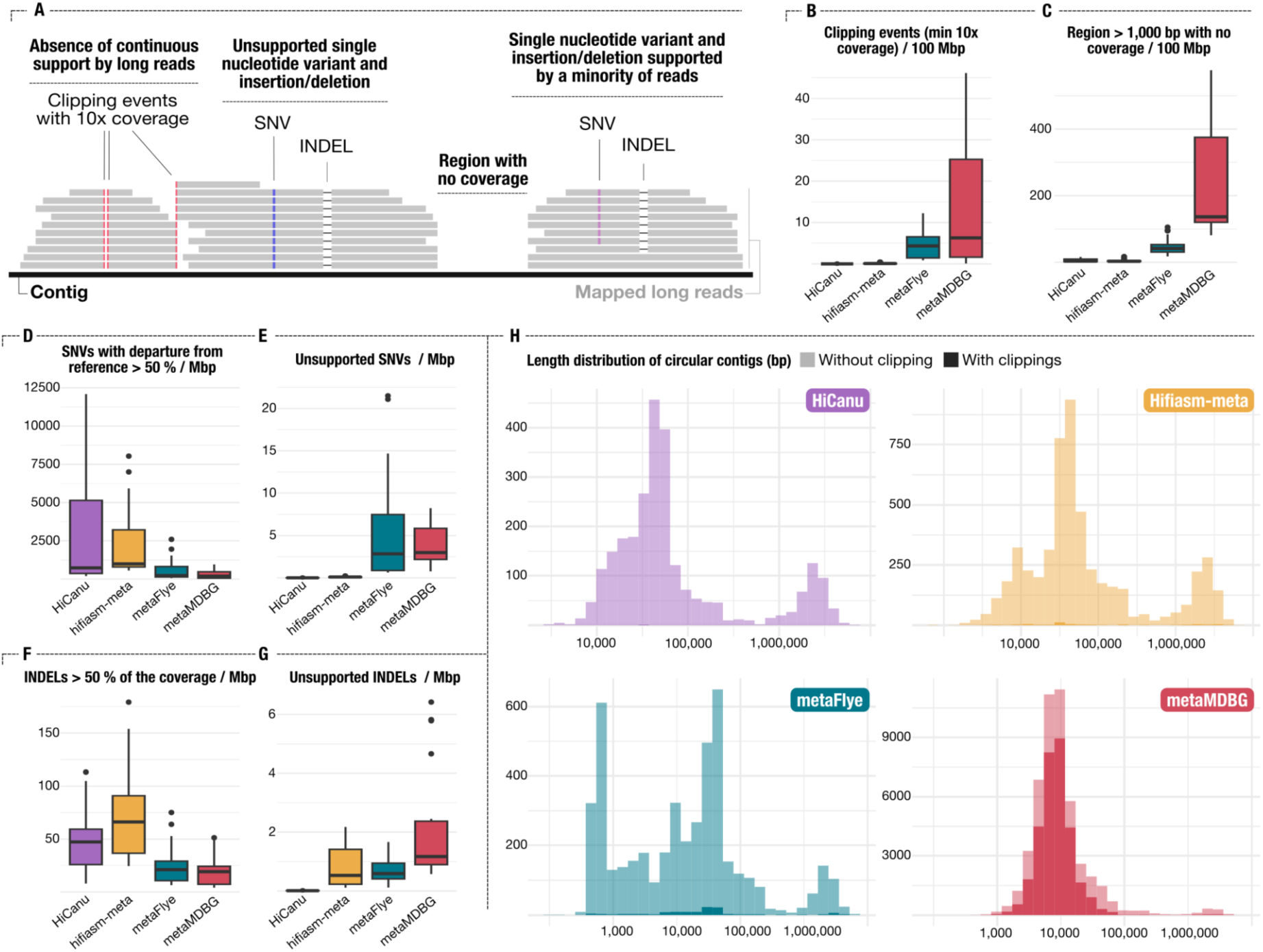
Assembly errors across assembler. (A) A schematic representation of long reads mapping to a contig with multiple types of read disagreement with the reference, including indel and single nucleotide variants representing more than half or all the coverage, and clipping events spanning the entire coverage. All metrics for B, C, D, E, F, G are normalized by assembly size and exclude the two mock community metagenomes. (B) Number of clipping events supported by at least 10 reads. (C) Number of regions over 1,000 bp with no apparent coverage. Number of single-nucleotide variants representing (D) > 50% or (E) all the coverage (G) at a given locus. Distribution of (F) indels > 50% of the coverage, or (G) all the coverage. (H) Length distribution of circular contigs by each assembler. The darker color represents the distribution of circular contigs with at least one clipping event.

The assembly metrics differed greatly between sample types and assembly algorithms. For instance, all human gut metagenomes were assembled to a similar total size by all assemblers, but not the surface ocean samples (Supplementary Table 2), for which metaMDBG produced 610% more assembled sequences compared to HiCanu on average. We found high confidence read clipping events (i.e., 100% clipping at locations with at least 10X coverage) in at least one sample from each assembler (Supplementary Table 2). However, the frequency of these events normalized by the assembly size showed that metaFlye and metaMDBG generated up to 180 times more clipping events compared HiCanu and hifiasm-meta for the same samples (Figure 1B). The number of clipping events were particularly high in the surface ocean samples, where assemblies from metaMDBG had over three orders of magnitude more clipping events compared to hifiasm-meta (Supplementary Table 2). Overall, clipping events affected up to 5.6% of contigs longer than 10,000 nucleotides reported by metaMDBG. We also computed the number of regions longer than 1,000 bp with no apparent coverage by individual long reads and found that the occurrence of regions that are not supported by any read were as pervasive as clipping events. While this issue also affected all assemblers (Figure 1C, Supplementary Table 2), metaMDBG led the pack with up to 5.3% of all contigs longer than 10,000 nucleotides with zero-coverage regions.

Reporting of contig circularity was a common feature to all assemblers, however, the number of circular contigs in final assemblies also varied between algorithms (Figure 1H, Supplementary Table 2). MetaMDBG generated substantially more circular contigs than the other assemblers, notably in surface ocean metagenomes. Overall, we found at least one clipping event for a large proportion of circular contigs reported by metaMDBG (Figure 1H), which, in some cases represented up to 77% of the circular contigs in a sample (e.g., the sample HADS 013, Supplementary Table 2).

In addition to the clipping events, we reported the frequency of single-nucleotide variants (SNVs) and insertion/deletion events (INDELs) in the assembled contigs that were either (1) represented in only a minority of long reads where the final assembly included a nucleotide or INDEL that did not match to the most frequent nucleotide or INDEL in long reads (Figure 1A, D, F), or (2) represented by none of the long reads, a more severe case where the final assembly included a nucleotide or INDEL that did not occur in any of the long reads matching that context (Figure 1A, E, F). After normalizing based on the assembly size, HiCanu and hifiasm-meta had the most SNVs and INDELs that represented only a minority of long reads at a given position, while metaFlye and metaMDBG had the most SNVs and INDELs that were not supported by any long reads. The latter case affected thousands of genes in all assemblies (Supplementary Table 2), leading to genes with incorrect amino acid sequences due to the impact of INDELs on open reading frames. Our detailed investigation of individual clipping events associated their occurrence with a few recurring classes of erroneous reporting of contigs, including chimeras, premature circularization, haplotyping issues, false duplications, and non-existent sequences, for which we offer an incomplete list of examples below.

### Chimeric contigs

Our inspection of contigs with a high proportion of read clipping events revealed chimeric contigs. In some cases, chimeras brought together sequences from taxa that belonged to distinct phyla (Figure 2). Most chimeras brought together sequences from two distinct taxa, but cases that merged sequences from more than three organisms were not uncommon (Figure 2), and they sometimes combined sequences from distinct domains of life. In one extreme case, metaMDBG formed a sequence that included regions from organisms that originated from Euryarchaeota, Pseudomonadota, Bacteriodota, and Cyanobacteria (Figure 2). We acknowledge that the list of contigs we surveyed here for chimerism is far from exhaustive, and furthermore the reliance of clipping events alone can occasionally miss chimeric contigs. For instance, we stumbled upon a suspiciously long contig. Even though this 7.38 Mbp contig was not flagged by the frequency of read clipping events, our manual inspection showed that it contained two sequence-discrete Lachnospiraceae populations metaMDBG reported from the sample Human O1 (Supplementary Figure Fig 1).

**Figure 2.**
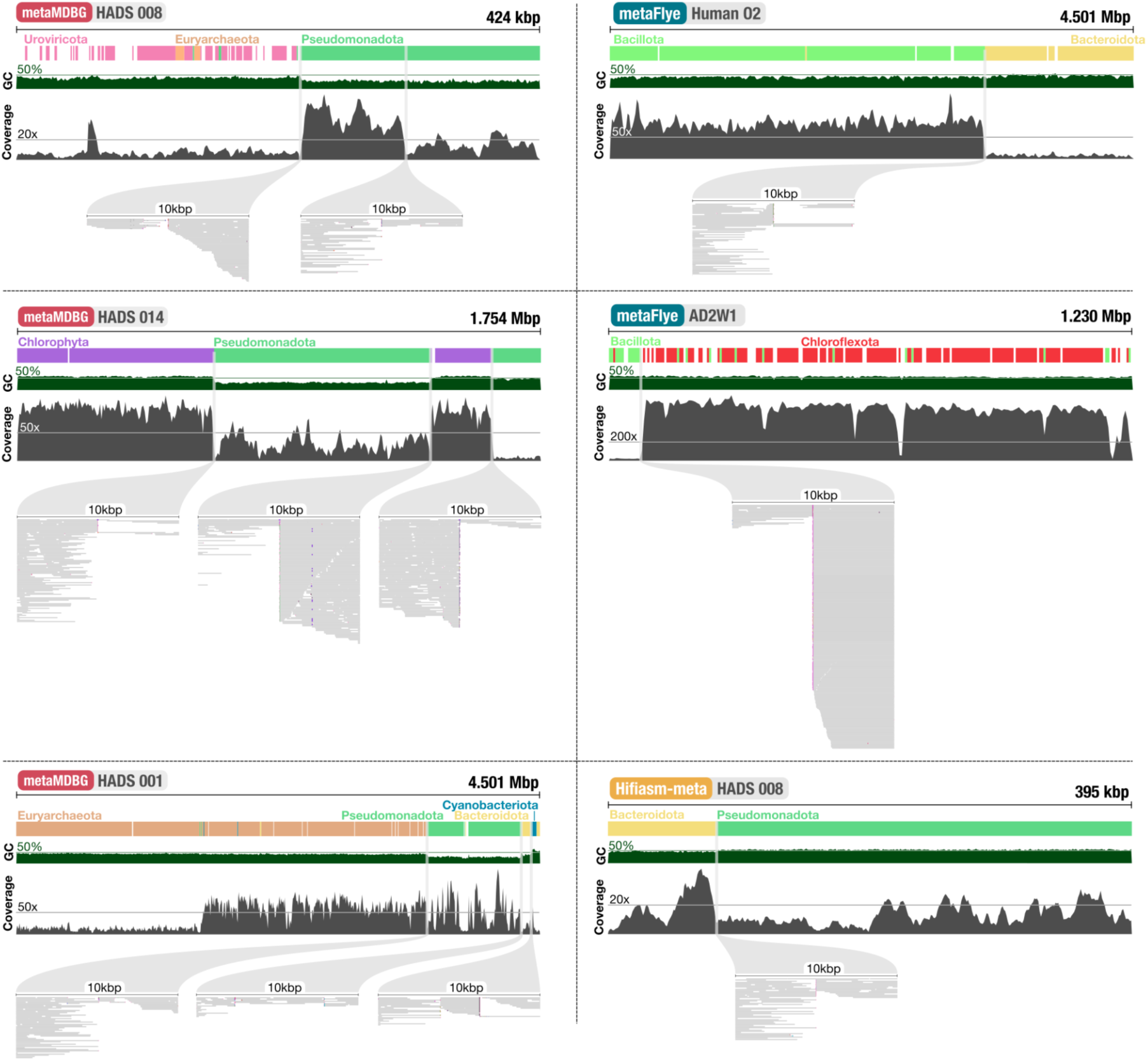
**Multi-domain and multi-phyla contigs**. Six contigs from metaMDBG, metaFlye and hifiasm-meta. For each contigs we displayed the GC content, coverage in the metagenomics reads used for their assembly, gene level taxonomy. For each assembly breakpoint, we display a zoomed-in detail of the read mapping from IGV. In these subplots, red arrows at the end of the mapped read indicate clipping and the coloring at the end of these reads indicates that the following portion of the read mapped to another contig and similar colors indicate that multiple reads continue to map on the same contig. Blue markers indicate large indels (> 150bp).

The highly dangerous nature of chimeric contigs for downstream analyses is dampened by the straightforward nature of their identification by anyone who carefully investigates their data: most chimeric contigs that erroneously connect genomic regions from two or more distinct populations can be easily identified using a series of genomic markers such as a sudden shift in GC content, read coverage, and gene-level taxonomy, or through unexpected inventories of single-copy core genes (SCGs). However, with the increasing tendency of researchers to generate large metagenome-derived genome compendiums (Pasolli et al. 2019; Parks et al. 2017; Ma et al. 2023; Nayfach et al. 2021; Almeida et al. 2021), such evaluations are rarely, if ever, conducted. In an ideal world, the burden of resolving chimeras should never fall on the shoulders of the end-users of assembly algorithms.

### Premature circularization

To deliver one of the most sought after promises of long-read sequencing, most long-read assemblers contain built-in features to circularize contigs and report potentially complete microbial and mobile genetic element genomes. Premature circularization, the reporting of a contig as circular when it omits parts of the genome it originates from, is a form of error with dangerous downstream implications. Yet, our reanalyses showed that the algorithmic features that reconstructed circular contigs in long-read assemblers, especially metaMDBG, and to a lesser degree hifhasm-meta, were far from reliable.

For instance, an archaeal genome that belongs to the genus *Methanothrix* recovered from the AD Sludge sample as a circular genome by hifiasm-meta represents a clear example of the nature and implications of premature circularization. Through an analysis with additional genomes from the NCBI’s RefSeq collection we confirmed that the circular genome was missing a large fraction of the core *Methanothrix* pangenome (Figure 3B, light blue), including key metabolic modules for methanogenesis that were common to all *Methanothrix* genomes (Figure 3C). In this case, early circularization occurred near a transposase (Figure 1D, Supplementary Figure 2). Nearly all the reads clipping on either side of the transposase had supplementary mapping to another contig in the assembly output (Figure 3B, C, medium blue) encoding the missing metabolic module. The combination of the circular genome and the additional contig matched the other genomes in the genus-level pangenome (Figure 3B, C, dark blue), suggesting that both contigs belonged to the same population, and neither of them were circular by themselves.

**Figure 3.**
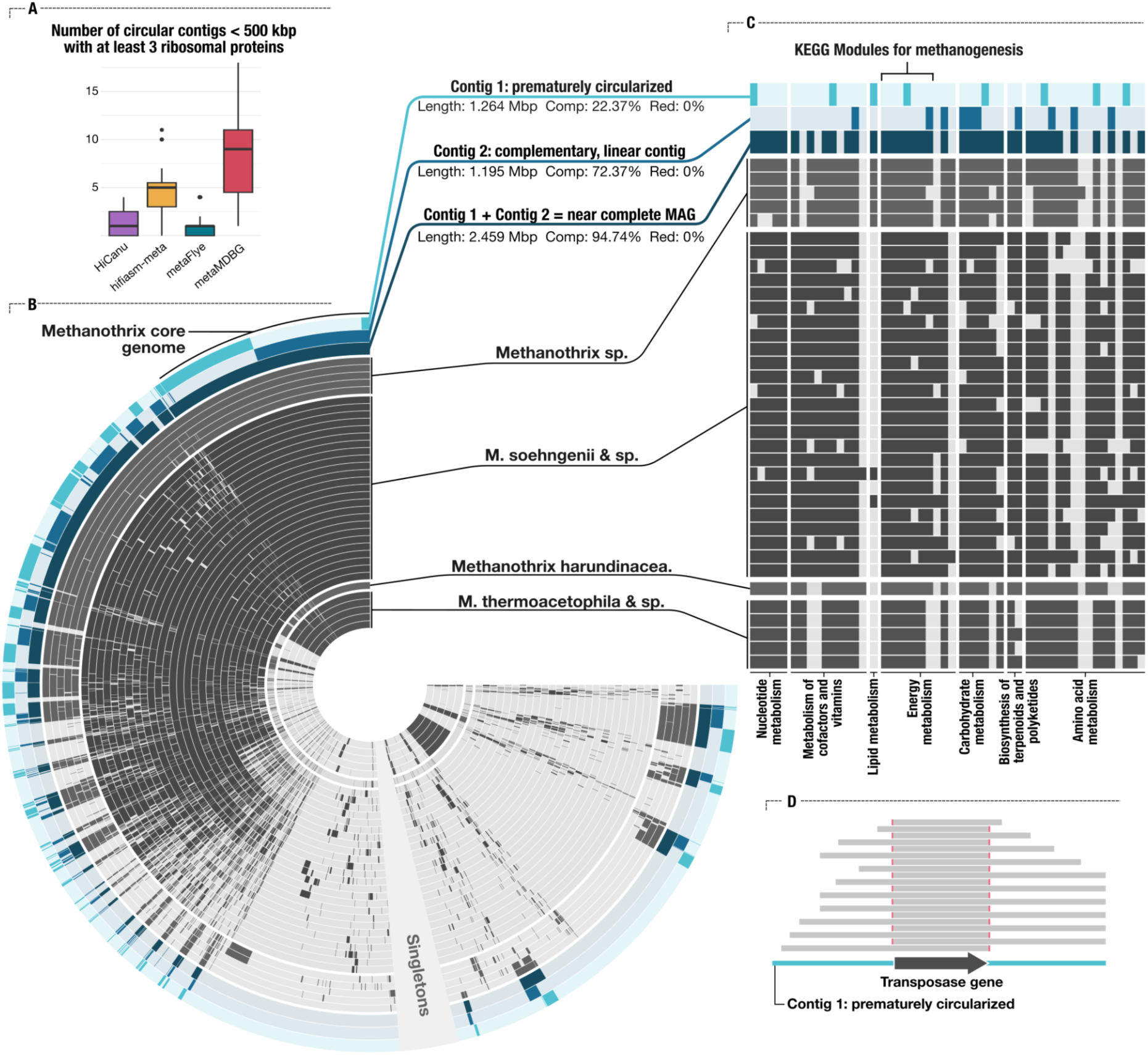
**Premature circularization of a *Methanothrix* genome**. (A) Frequency of circular contigs under 500kbp with a minimum of 3 ribosomal proteins. Each point represents one assembly. (B) A pangenomic analysis of all publicly available *Methanothrix* genomes from the NCBI’s RefSeq database completed with the so-called circular genome of *Methanothrix* assembled from the sample AD Sludge by hifiasm-meta (light blue), as well as a contig from the same assembly which correspond to the rest of the missing Methanothrix genome (medium blue) and the combination of these two contigs (dark blue). (C) KEGG metabolic module completion of all genomes and contigs in (B). (D) A schematic representation of the reads mapping over a transposase gene in the prematurely circularized contigs (light blue in B and C) showing the lack of reads support around the gene, the full figure is available in Supplementary Figure 2.

Comparing the overall accuracy of circularization across assemblers is difficult. For instance, low completion estimates based on single-copy core genes can serve as a quick filter, but not all prematurely circularized genomes will have low completion estimates since missing genomic content will not always contain single-copy core genes. Length comparisons between circular contigs and known genomes in databases could offer another means of scrutiny, but this strategy will not be effective against poorly studied clades, or those that have no representation in public databases. Nevertheless, here we conservatively assumed that circular contigs that were under 500 kbp and contained at least 3 ribosomal proteins were most likely circularized erroneously, and we further assumed that the rates at which they are generated by each assembler could serve as a proxy for an overall estimate of the rate of premature circularization events across assemblers. For this range, metaMDBG reported about twice as many circular contigs than hifiasm-meta, and about four times more circular contigs than HiCanu and metaFlye (Figure 3A). Assuming that easily identifiable events of premature circularization errors are reasonable proxies for the rate of all premature circularization errors, this result suggests that metaMDBG and hifiasm-meta are more prone to premature circularization errors compared to other assemblers, such as HiCanu and metaFlye. Interestingly, our estimates of the rate of false circularization come very close to the benchmarking results shared by the authors in the original metaMDBG publication (Benoit et al. 2024), in which they report two times more circular contigs than hifiasm-meta and four times more circular contigs than metaFlye from the human gut. While Benoit et al. present the high rate of circularization as a strength of metaMDBG in comparison to other assemblers (Benoit et al. 2024), our observations suggest that the higher rate of circularization may be a result of a higher tendency to report non-circular elements as circular (Figure 1, Figure 3, Supplementary Figure 2, Supplementary Table 2).

Given the vast difference in quality and perception between a draft genome and a circular genome, it is essential for assembly algorithms that promise circular genomes to be conservative in their decision making. The identification of prematurely circularized contigs in assembly results will be a notoriously difficult task for end-users, especially when the missing genomic context is relatively short, or circular contigs represent plasmids or viruses. Thus, it would be ideal if the circular sequences are validated more rigorously by the assembler prior to the final reporting.

### Haplotyping errors, false duplications, and non-existent sequences

Accurate reconstruction of genomic variation is essential to associate within-population structural differences to ecological or evolutionary phenotypes. However, resolving genomic regions that differ between otherwise very closely related subpopulations is a major challenge for de novo assemblers (Liao et al. 2019). Assemblers may resolve such structural complexity (1) by reporting separate contigs for variable sequences and conserved regions that flank them, (2) by reporting the most prevalent variable region along with the flanking conserved regions in a single contig and alternative regions in shorter contigs, or (3) by duplicating conserved flanking regions in multiple contigs that describe each of the variable region and their surrounding as separate contigs in a haplotype-aware fashion. Our survey of long-read assembly results revealed unexpected haplotyping decisions in multiple recurrent forms. In some cases, assemblers concatenated subpopulation-specific variable regions flanked by conserved loci, rather than reporting only one of them accurately (Figure 4A). As a result, long-reads that map to these regions are clipped at the end of their respective sub-population sequence. In other cases, the final contig represented a variable region found in a minor subpopulation supported by a small number of long-reads or a single one (Figure 4B), violating the logical expectation to recover a consensus sequence that represents the most abundant sub-population.

**Figure 4.**
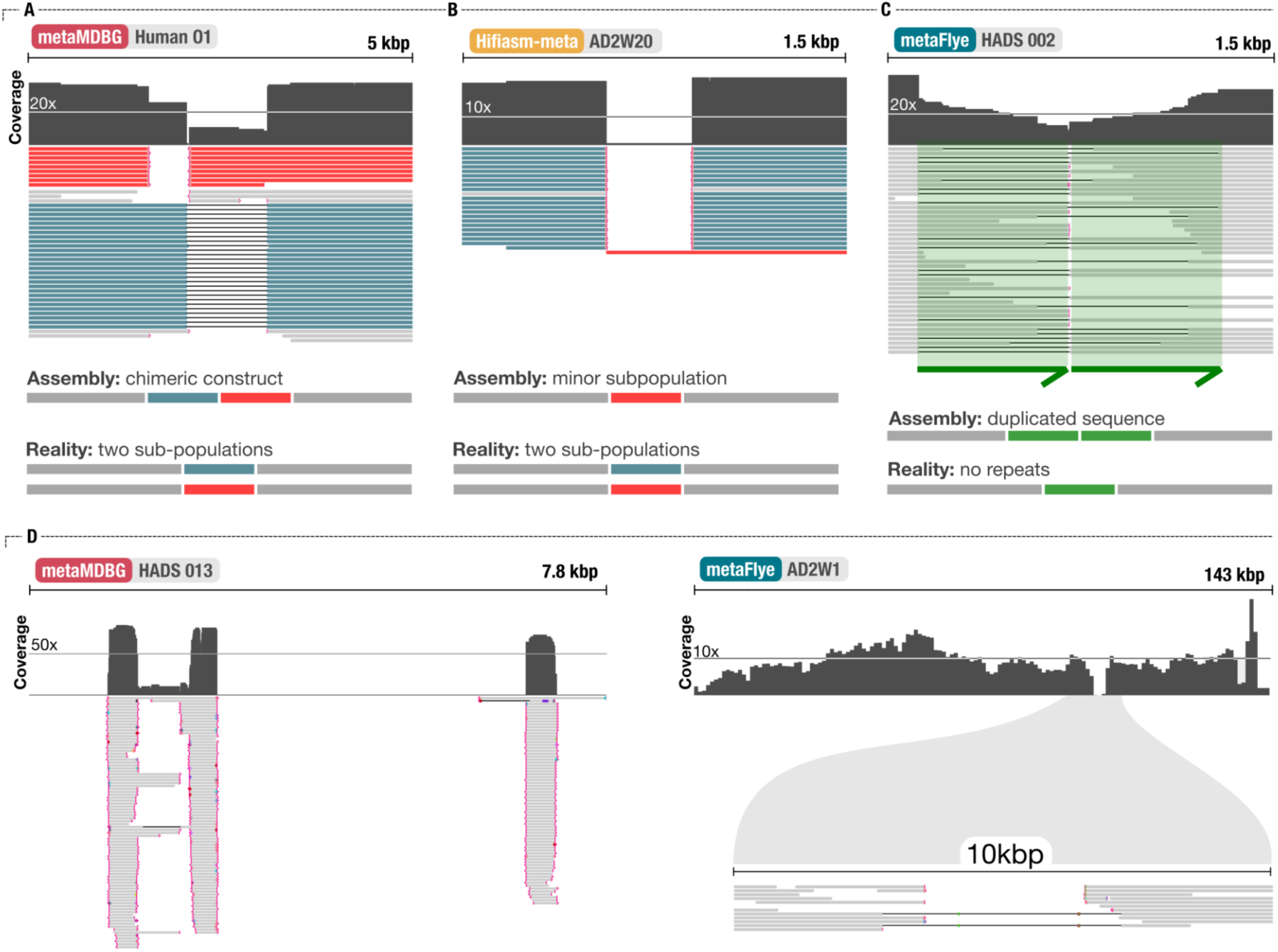
**Prototypical mapping artifacts and their putative origin**. (A) A chimeric sequence assembled from two subpopulations. At a conserved locus, two subpopulations existed with their own and distinct sequence. The assembled contig contains all or a part of each subpopulation specific sequence resulting in a chimeric construct. (B) Another example of a variable genomic site, but in this example the contig sequence contains the sequence of a very minor subpopulation, supported by only one read. (C) Duplicated sequence only found in the assembly, not supported by any long reads. (D) Two contigs assembled from metaMDBG (left) and metaFlye (left) presenting large regions with no coverage. We blasted these regions back to the long reads and found no hits. Coverage visualization was exported from the anvi’o interactive interface (left) or the IGV software (right) and the read mapping visualization was from IGV as well. Indel smaller than 150bp as well as mismatches are not displayed in the mapping. Red markers at the end of reads indicate read clipping.

We also identified cases where assemblers reported false genomic duplications in assembled contigs that were not supported by any long-read. Such false repeats were often manifested by high-frequency read clipping events and appeared as long direct repeats that had low likelihood to be present in the target genomic context due to the sudden decrease in read coverage and/or massive inserts in mapped reads (Figure 4C). Yet another anomaly was the reporting of sequences that did not exist. BLAST-searching zero-coverage regions from contigs that are longer than 500 bp against the database of raw long reads (Supplementary Table 3), we confirmed that the assembly outputs by metaMDBG and metaFlye occasionally included up to over 5,000 bp long regions that have no homology to any of the input long reads (Figure 4D). False duplications and reporting of non-existent sequences in assembly results are unexpected behaviours from any assembler and can lead to spurious open reading frames or the omission of genuine ones.

### Excessive repeats

A recurrent and puzzling observation throughout our manual inspections of assembly results was the astonishing number of repeats that occasionally made up the entirety of some contigs. Yet, these repeats were not caught by our survey of read clipping event frequencies to mark regions of concern as repeats rarely resulted in 100% read clipping events to pass our filter. Thus, we characterized repeats without any read mapping data but by aligning each contig to itself. We marked any region of a contig as a ‘repeat’ if it was longer than 200 bp and occurred multiple times in the same contig with at least 80% identity. Our survey of the frequency and distribution of such repeats across contigs under 50 kbp showed that each assembler reported contigs with repeats. Since repeats are common in nature, and the improved ability to resolve repeats is one of the strengths of long-read sequencing, this finding alone is not concerning. However, the nature of repeats in assemblies revealed by dot plots often displayed surprisingly intricate patterns suggested a high likelihood that they were assembly artefacts, rather than natural genomic organisations (Figure 5A). While naturally occurring tandem repeats, inversions, or palindromes could lead to similar dot plots, our manual inspection of individual cases revealed a variety of errors that spanned from duplicated reporting of known circular plasmids to contigs with multiple repeats that are not supported by any long read.

**Figure 5.**
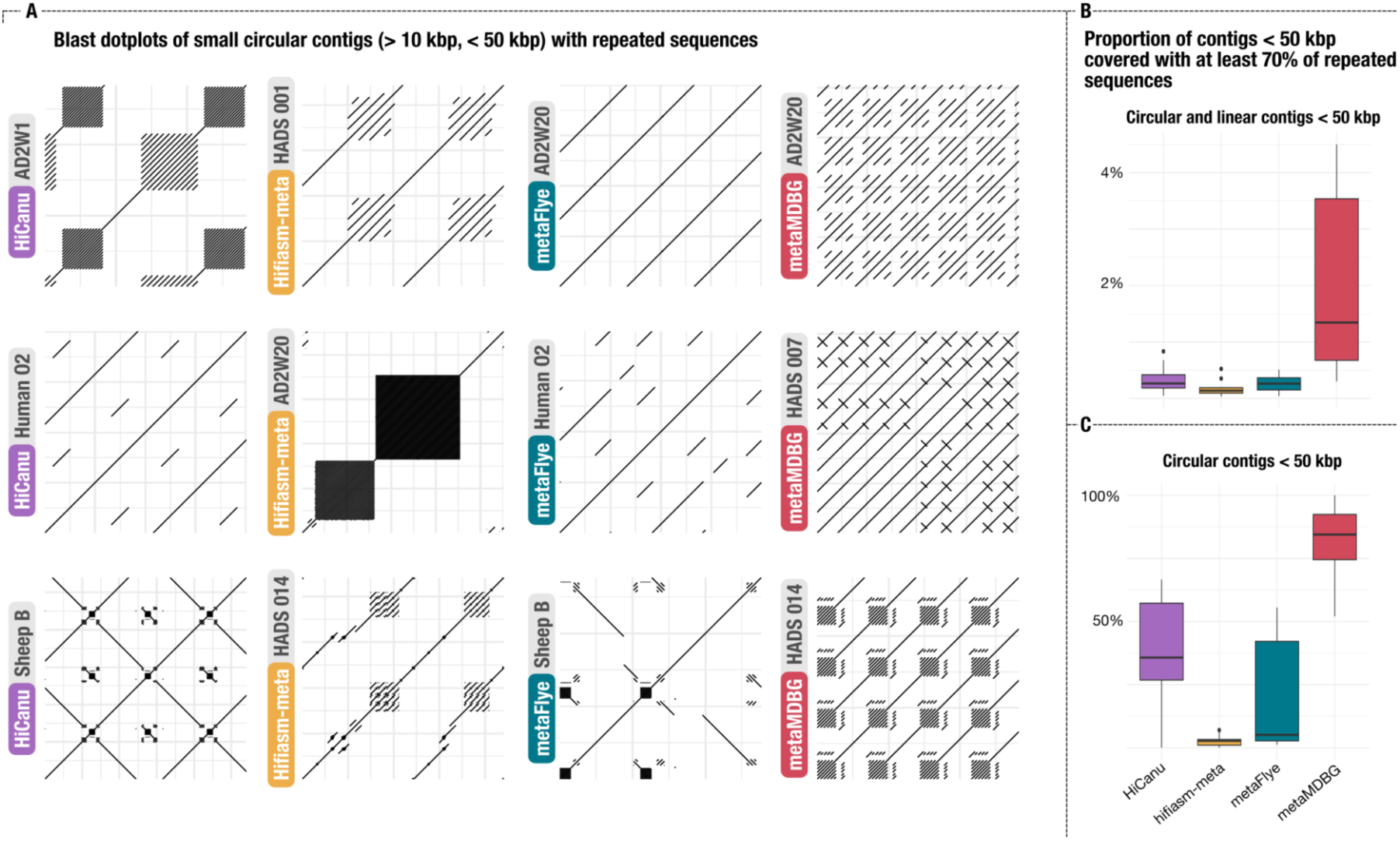
Repeats across short contigs. (A) Dotplots of reciprocal blast results from 12 short circular contigs showing high amounts of repeated sequences. (B,C) Proportion of contigs with at least 70% of their length duplicated at least once. Result computed for all short contigs under 50kbp (B) and only for circular contigs under 50kbp (C).

Repeats differed in their length, identity, and frequency across algorithms. The per sample average length and sequence identity of repeats varied between 600 bp to over 4,000 bp, and 88% to 96%, respectively. MetaMDBG generated more repeats than any other assembler (Figure 5B), up to over 300,000 repeats in a single assembly (Supplementary Table 2), exceeding the number of repeats reported by HiCanu for the same sample 280 times. The average length of repeats were relatively short, yet we found repeats that were up to 40,000 nucleotides long that were due to multiple tandem repeats forming near contig-long and overlapping repeats, and repeats occurred as much as over a 1,000 times in a single contig (Supplementary Table 2). To summarize the number and proportion of contigs reported by each assembler with an excessive number of repeats, we conservatively searched for assembled sequences in which at least 70% of the sequence was composed of repeats. This search revealed that metaMDBG generated on average 2.8% of contigs under 50,000 bp with a high amount of repeats, a rate that was over 6 times higher than its runner up, HiCanu (Figure 5B). When we limited our survey to circular contigs, the proportion of repeat-rich contigs skyrocketed across all sample types for all assemblers, with the exception of hifiasm-meta (Figure 5C, Supplementary Table 2). For instance, 87% of all circular contigs under 50 kbp reported by metaMDBG were largely composed of repeats, and in some samples, such as Chicken, this number reached to 100%, suggesting that artifactual repeats represent at least one of the factors that lead to false circularization (Figure 5C). Marine metagenomes were particularly difficult for all assemblers. In this sample type metaMDBG reported the highest number of circular contigs (Supplementary Table 2), however, 92% of them under 50,000 bp were composed of repeats (Supplementary Table 2). While metaFlye performed as well as hifiasm-meta in most other sample types, up to 55% of circular contigs metaFlye generated from marine metagenomes were largely composed of repeats (Supplementary Table 2).

These results show that the exciting prospects of recovering complete and circular plasmid and virus genomes from long-read sequencing of complex metagenomes is still far from being realized, and that the state-of-the-art long-read assemblers often fall short when handling short circular elements.

### Mock datasets are useful, but can yield misleading insights into the accuracy of algorithms in real-world applications

Popular mock datasets such as Zymo-HiFi D6331 and ATCC MSA-1003 are commonly used for benchmarking long-read assemblers. While their known composition constitutes a reasonable starting point, the mock datasets do not represent the complexity of the natural samples, as also noted by the authors of metaFlye (Kolmogorov et al. 2020). Their utility to test assemblers is further reduced when benchmarks that employ mock communities simply align contigs that emerge from the assemblies to reference genomes without comprehensive reporting of other assembly metrics (Benoit et al. 2024; Kolmogorov et al. 2020; Feng et al. 2022). For instance, while hifiasm-meta reports favorable outcomes given the reference genomes in mock datasets (Feng et al. 2022), in our tests the algorithm generated massive assemblies for each mock dataset, where the final assembly was 270 Mbp instead of the expected size of 93 Mbp for the Zymo-HiFi D6331, and it was 948 Mbp instead of the expected size of 66.44 Mbp for the ATCC MSA-1003, resulting in the lowest N50 values and the highest number of clipping events across all assemblers. Given its performance with the mock datasets, one may expect hifiasm-meta to perform poorly in its applications to complex metagenomes. Yet, in marine samples, hifiasm-meta was first in N50, and second to metaMDBG in assembly size with two orders of magnitude less clipping errors, suggesting that the performance of an assembler with mock communities may not predict its performance with real-world datasets.

While mock communities do not represent the diversity and complexity found in real-world samples, they shine in one fundamental way: the known genomic makeup of input organisms to identify glaring issues post-assembly. The Zymo-HiFi D6331 includes five different strains of *E. coli*, which yields sequencing data with complex cases for assemblers due to the presence of highly conserved and divergent genomic regions. HiCanu and hifiasm-meta were both successful as reconstructing at least one of the five *E. coli* genomes (Supplementary Figure 3), while metaMDBG reported a circular contig that corresponded to a chimeric genome (Figure 6). Based on pairwise ANI comparisons, this genome appeared to be most similar to B1109, one of the *E. coli* strains in the Zymo-HiFi D6331 mock dataset (Figure 5B, Supplementary Table 4). However, the pangenome of the five *E. coli* genomes and the metaMDBG circular contig showed that not only it was missing a portion of the B1109 genome, but also including genomic regions exclusive to other *E. coli* strains and absent in B1109 (Figure 6A). Clipping events also captured chimeric regions and revealed a ∼10 kbp locus with no coverage (Figure 6C, D). That region without coverage was duplicated in two other contigs, suggesting that long reads were preferentially recruited there (Supplementary Table 5). Using ANI values that are calculated from local alignments of assembled contigs from mock datasets to the collection of reference genomes they contain risks masking glaring issues such as phantom sequences, chimeras, and unexpectedly large assembly results, and invalidates perhaps the only useful aspect of mock datasets while inflating the accuracy of assembly algorithms.

**Figure 6.**
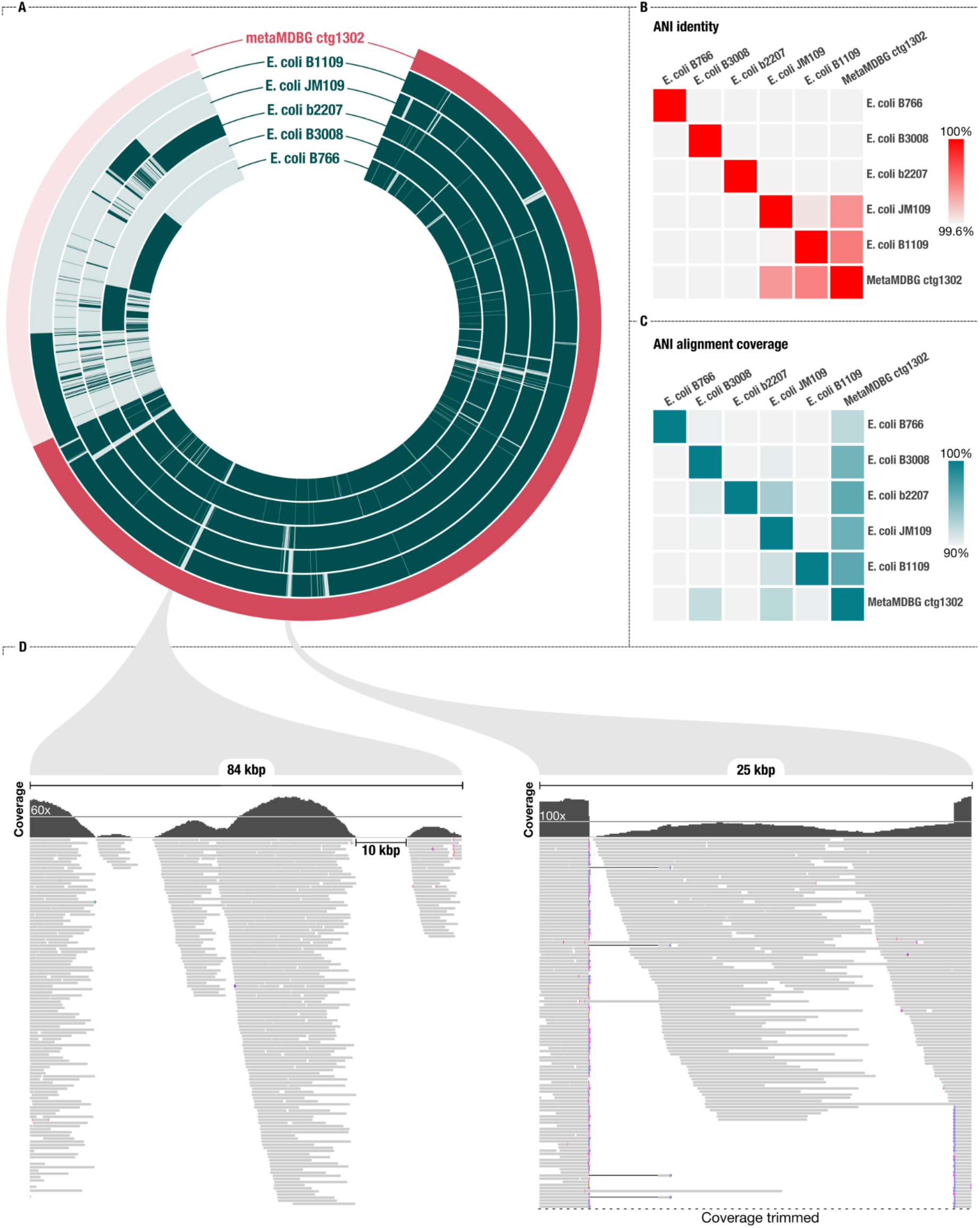
**Assembled E. coli from a mock community does not match reference genomes**. (A) A pangenome analysis of the *E. coli* reference genome used in the Zymo mock community and the *E. coli* circular genome assembled by metaMDBG. Each layer corresponds to a genome and the coloring represents the presence/absence of a gene cluster from each genome. The gene clusters are organized by the synteny of genes in the metaMDBG contig. (B) Genome pairwise average nucleotide identity and (C) alignment coverage. (D) Details of the long read mapping over two genomic regions highlighted in the first panel. These regions included genes not found in E. coli B1109 (closest reference genome by ANI metrics), but rather found in other E. coli genomes.

## Conclusions

Our findings highlight the need for rigorous evaluation of long-read assembly algorithms beyond benchmarks that typically prioritize runtime, contig length, or the number of circular contigs. We demonstrate that identifying the most terrifying assembly errors can be as simple as mapping individual reads back to assembled contigs. Some popular long-read assemblers like metaFlye include a correction step that uses the long reads to polish the assembly, and could leverage mapping information like clipping events to generate more accurate assemblies. We believe assembly algorithms should provide better options for error correction using input reads, rather than offloading this burden onto end users who may lack the time, expertise, or computational resources to perform such refinements themselves. The biotechnology, biomedical, and basic research communities rely on high-quality assemblies, and trustworthy results are critical for progress in life sciences as well as the quality of public genome databases. Given the critical role of assembly in genomics, most researchers will likely prefer longer runtimes or fewer circular contigs over algorithms that are fast or produce erroneous sequences.

## Materials and methods

The URL https://merenlab.org/data/benchmarking-long-read-assemblers/ serves our bioinformatics workflow to reproduce our findings or apply the same approaches to evaluate additional assemblers or datasets.

### Datasets

We downloaded a set of publicly available HiFi PacBio metagenomes matching the ones used in long-read assembler publications. To complete and expand the set of biomes, we included eight surface ocean metagenomes. Supplementary Table 1 includes a comprehensive description of these data and their accession numbers.

### Assembly algorithms

We used 4 different assemblers: HiCanu v2.2 (Nurk et al. 2020), hifiasm-meta v0.3 (Feng et al. 2022), metaFlye v2.9.5 and metaMDBG v0.3, v1 and v1.1 (Benoit et al. 2024). For HiCanu, we used the parameters maxInputCoverage=1000 genomeSize=100m batMemory=200 and-pacbio-hifi. We used hifiasm-meta with the default parameters. For metaFlye, we used the parameters--meta--pacbio-hifi. We included three versions of metaMDBG; v0.3, which is the original published version of the software, and v1 which was released August 2024, and v1.1 released in December 2025. The results for v0.3 and v.1.1 can be found in the supplementary material while the results from v1 are shown in this manuscript. For metaMDBG v1 and v1.1, we used the parameters--in-hifi.

### Read mapping

We mapped the metagenomics long-reads back to their respective assembly by the four assemblers using minimap2 v2.28 (Li 2018) with the following parameters:-ax map-hifi-p1 -- secondary-seq. This set of parameters allows secondary mapping when the alignment score is as good as the primary mapping score, i.e. multi mapping; as well as to keep the sequence for secondary mapping in the output files so that secondary mapping is properly considered in downstream analyses. We processed the resulting alimgent files using samtools v1.17 (Li et al. 2009).

### Processing of assembled contigs and read mapping results

We used anvi’o development branch of v8.1 (Eren et al. 2021) to generate contigs databases for each assembly with the command ‘anvi-gen-contigs-database’, which performed gene calls using prodigal v2.6.3 (Hyatt et al. 2010). The anvi’o programs ‘anvi-run-ncbi-cogs’ and ‘anvi-run-kegg-kofams’ annotated genes with function suing the NCBI’s Clusters of Orthologous Genes (COGs) (Galperin et al. 2015) database and KEGG KOfams (Aramaki et al. 2020), respectively, while ‘anvi-run-hmms’ identified single-copy core genes in these sequences which we associated with taxonomy data from the GTDB (Parks et al. 2022) using ‘anvi-run-scg-taxonomy’. We estimated the completeness of KEGG KOfam modules using ‘anvi-estimate-metabolism’. We finally used ‘anvi-profile’ to process the read mapping data to recover coverage values, single-nucleotide variants (SNVs) and insertion/deletion events (INDELs).

### Pangenomic analyses

To compute a pangenome for *Methanothrix* we first acquired publically available genomes from NCBI’s RefSeq database using the program ‘ncbi-genome-download’ (available from https://github.com/kblin/ncbi-genome-download) with the parameters “--assembly-level all -- genera Methanothrix”. For the *E. coli* pangenome, we downloaded the original genomes used to create the mock dataset: https://s3.amazonaws.com/zymo-files/BioPool/D6331.refseq.zip. For both pangenomics analysis, we used the program ‘anvi-run-workflow’ to run the anvi’o pangenomics workflow implemented in Snakemake (Köster and Rahmann 2012), which used DIMAOND v2.1.8 (Buchfink, Xie, and Huson 2015) to identify gene clusters as described previously (Delmont and Eren 2018). We used the program ‘anvi-display-pan’ to visualize and summarize the pangenomes.

### Identification of assembly errors

To identify potential assembly errors based on clipping events, we developed a program within the anvi’o platform, ‘anvi-script-find-misassembly’ (https://anvio.org/m/anvi-script-find-misassemblies), which takes a single BAM file of long-read mapping results. The script searches for premature end of alignments, i.e., clipping events, and reports positions where a proportion of reads that are clipped exceeds a user-defined threshold. The script also reports regions in contigs with no coverage. In addition, we used BLAST v2.16.0 (Camacho et al. 2009) to identify contigs covered by at least 70% of repeated sequences. We blasted each contig against itself using blastn default parameters, excluded the perfect reciprocal hit and transformed the remaining hits to a bed file and we used bedtools v2.31.1 (Quinlan and Hall 2010) to compute the breath of coverage. We computed additional statistics using a python script available in the link to the reproducible workflow

### Manual inspection of mapping results

We used IGV v2.17.4 (Robinson et al. 2017) and the anvi’o interactive interface to manually inspect genomic regions of interest and generate figures.

## Data availability

All metagenomes used in our study are publicly available through the NCBI, and Supplementary Table 1 lists their accession IDs. DOIs for intermediate data products are available at https://merenlab.org/data/benchmarking-long-read-assemblers/

## Acknowledgements

Authors recognize that developing assembly algorithms, especially for metagenomes, is a notoriously complex and difficult task, and they would like to acknowledge their deep appreciation of those who invest their time and skills in creating and maintaining them.

## Author contributions

FT, RS, JFB, and AME conceptualized the study. FT curated data, performed formal analyses, and developed software tools. FT, RS, JFB, and AME analyzed data and interpreted findings. FT and AME wrote the original draft of the study, all authors commented on and made suggestions, and approved the final manuscript.

## Ethics declarations

## Supplementary Information

### Supplementary Tables

*Supplementary Tables are also available via* doi:10.6084/m9.figshare.28831904.

**Supplementary Table 1**: Sample information and metagenome accession numbers.

**Supplementary Table 2:** Assembly metrics. (a) General statistics about the assemblies. (b) Long-read clipping across samples and assemblers. (c) Metrics about circular contigs, including clipping events. (d) SNVs and INDELs that are either supported by a minority of long reads (called’partial SNVs/INDELs’) or not supported by any long reads. (e) Prematurely circularized contigs.

Number of contigs less than 500 kbp with at least three ribosomal protein genes. (f) Frequency and metrics of repeats found in short contigs (less than 50 kbp).

**Supplementary Table 3:** Region of contigs with no apparent coverage and with no similarity (blast) with any long-reads.

**Supplementary Table 4:** Average Nucleotide Identity values for *E. coli* genomes. (a) ANI - Percent Identity. (b) ANI - Alignment coverage.

**Supplementary Table 5:** BLAST result of the *E. coli* sequence with no apparent coverage reported by the metaMDBG to the rest of the assembly.

### Supplementary Figures

*High-resolution figures are available via* doi:10.6084/m9.figshare.28831904.

---

**Figure S1:**
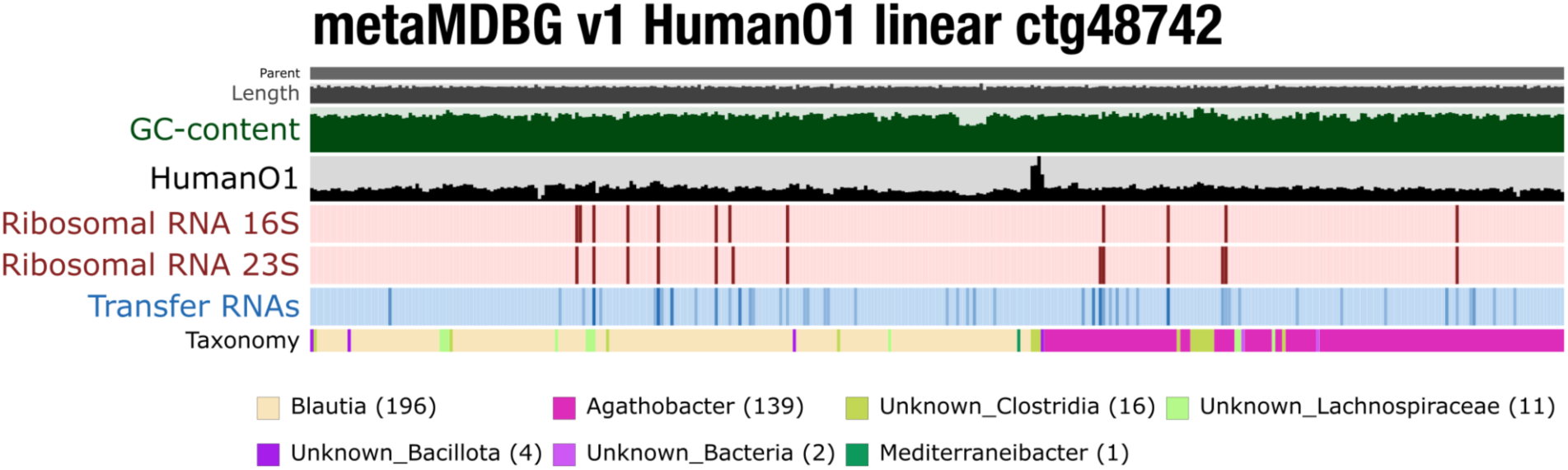
anvi’o interactive interface of a chimeric contig with no apparent clipping issues. The contig is 7.38 Mbp long and the estimated completion/redundancy are both 100%. Gene level taxonomy shows that two populations of Blautia and Agathobacter were merged into a single contig.

**Figure S2:**
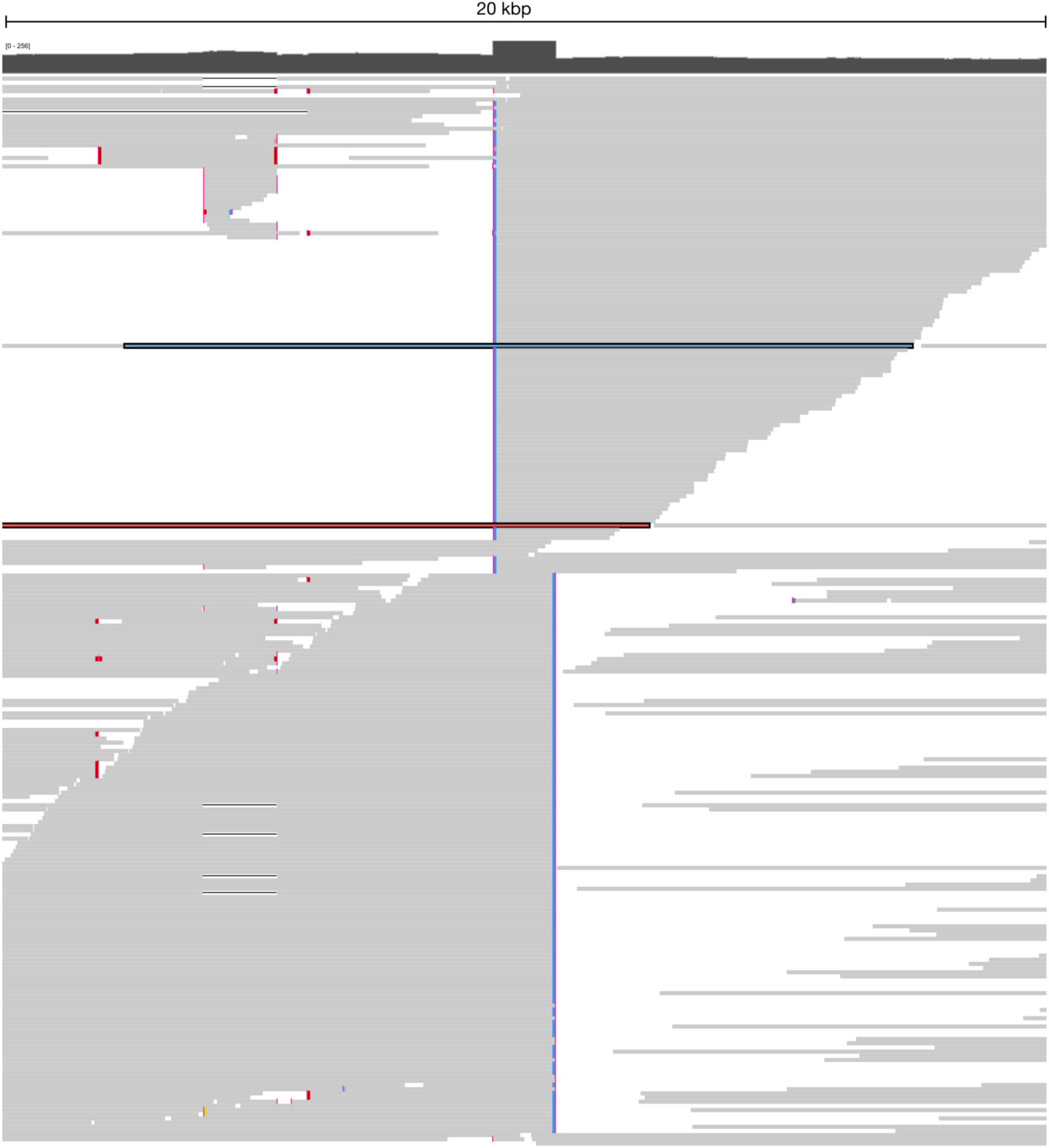
Long-read mapping of a 20kbp region in the prematurely circularized *Methanothrix* genome from Figure 2. The long-read overlap in the middle of the figure over a genomic region corresponding to a transposase gene. All long-read (except two, highlighted in red and blue),

**Figure S3:**
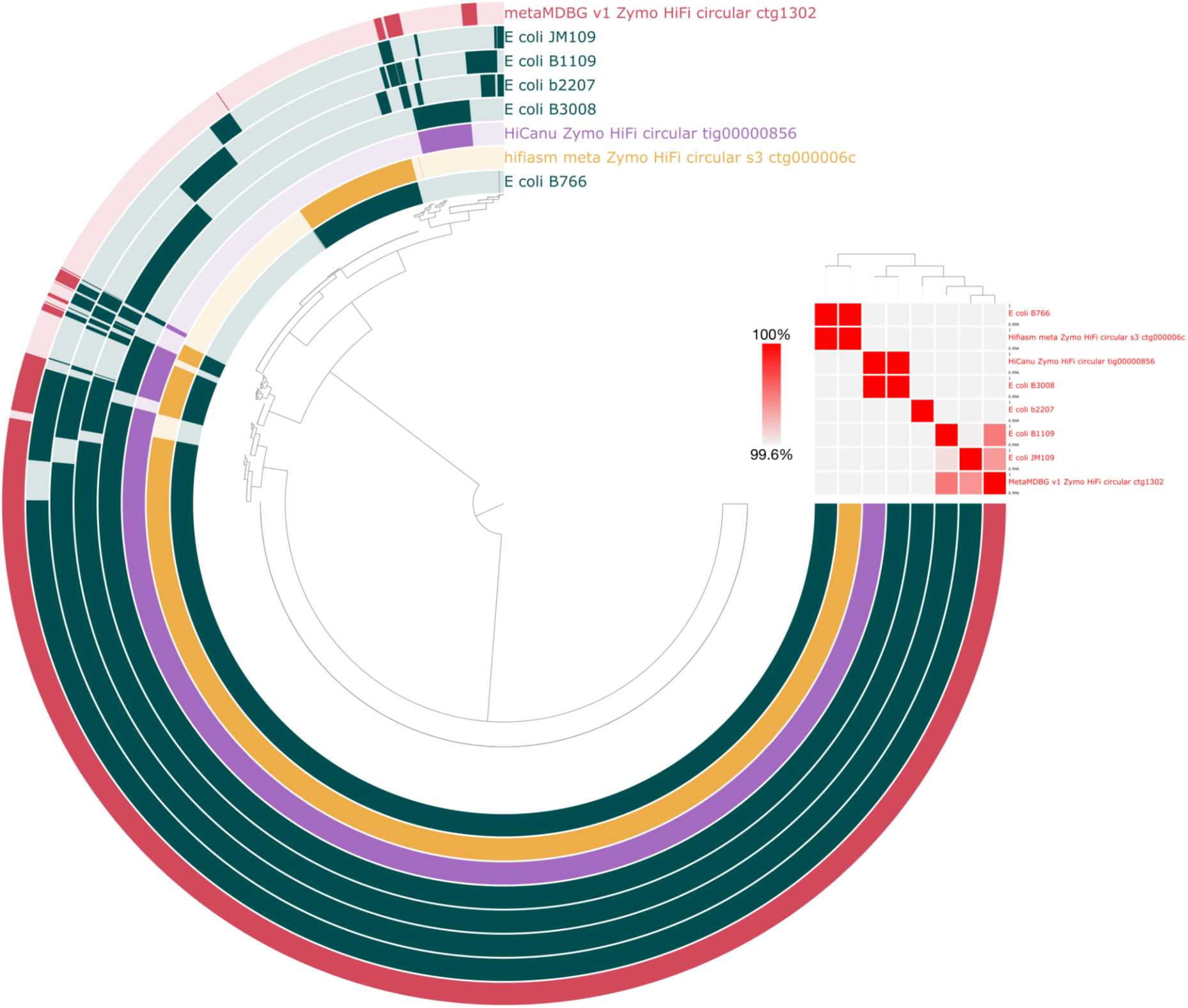
Pangenome analysis of the five *E. coli* genomes from the Zymo-HiFi D6331 mock community and the circular contigs generated by HiCanu, hifiasm-meta and metaMDBG matching to *E. coli.* Both circular contigs generated by HiCanu and hifiasm-meta were perfectly matching one of the five *E. coli* genomes while the contig from metaMDBG contained genes coming from multiple *E. coli* genomes.

